# Mucin anchoring of SARS-CoV-2 neutralizing nanobinders increases intranasal antiviral efficacy

**DOI:** 10.1101/2025.04.10.648177

**Authors:** Joelton Rocha Gomes, Juliette Jacquelin, Amélie Donchet, Nathalie Lejal, Quentin Nevers, Bernard Klonjkowski, Sophie Le Poder, Bernard Delmas, Nicolas Meunier

## Abstract

Respiratory viruses represent a major threat for human health, and despite effectiveness to limit the severity of the disease, they fail to block transmission. Recent advances have shown that the nasal cavity is the primal infection site of respiratory viruses, and thus represents a very attractive target for prophylactic treatment with antivirals in order to limit virus dissemination. However, mucociliary clearance limits the efficiency of a local treatment in the nasal cavity. We have previously developed a potent anti-Spike nanobinder blocking SARS-CoV-2 entry. Here, it was used as a proof of concept to show the strength of anchoring this synthetic antiviral to the mucin layer with a mucin-binding domain, increasing its residence time in the nasal cavity up to 6 hours post-instillation. This was very effective as a prophylactic treatment to limit infection of sentinel hamsters. Our strategy could be extended to antivirals against other major respiratory viruses such as RSV or influenza viruses, but also in other diseases by targeting specific epithelia increasing residence time and local concentration of the drug.

## Introduction

With nearly 25 million estimated deaths worldwide in less than four years, the COVID-19 pandemic demonstrated the necessity to better understand and combat the spread and transmission of respiratory viruses. The early infection of the nasal cavity suggests a major implication of the upper airways in the initiation and progression of the disease towards the lungs, a feature that has been confirmed experimentally in animal models for numerous respiratory viruses such as respiratory syncytial virus (RSV) and influenza viruses (Richard *et al*, 2020; Bryche *et al*, 2020; Hou *et al*, 2020). Blocking respiratory virus multiplication with antivirals delivered in the nose may therefore offer therapeutic benefit and prophylactic protection (Kozlov, 2022). Such a strategy would make treatment easier, allowing deposition of local drugs in higher concentration than with systemic treatment. Series of human neutralizing monoclonal IgG antibodies and nanobodies/VHH targeting respiratory viruses have been tested for systemic treatments with moderate success as they may not achieve therapeutic dose in the respiratory tract (Valdez-Cruz *et al*, 2021). Moreover, their delivery into the nose as a spray/aerosol may not be suitable, due to their cost and low stability (Mayor *et al*, 2021).

We previously produced an artificial protein (called C2; 158 amino acids long) targeting the receptor-binding domain of the SARS-CoV-2 Spike and exhibiting a very potent antiviral effect (Thébault *et al*, 2022). It displays an affinity for the Spike protein in the nanomolar range and neutralizes different variants of SARS-CoV-2 *in vitro*. However, an improved version of C2 only allows partial protection upon nasal instillation in a hamster model despite its high neutralization activity, due to low residence time in the nasal cavity. Indeed, the efficiency of mucociliary clearance in the nasal cavity challenges local treatment with drugs (Gizurarson, 2015). In the nose, specialized epithelial cells produce mucus as a ∼15 µm thin apical layer. This is mainly composed of a mucin network (3%) in water (95%) with two layers: the upper one thought to be cleared in ∼15 min and the inner defined by membrane-anchored mucins (Chen & Dulfano, 1978; Marttin *et al*, 1998). Mucins constitute a large family of heavily O-glycosylated proteins, defined by their variable tandem repeat (TR) domains, that cover all mucosal surfaces (Corfield, 2015). In the nasal cavity, Muc5b is mainly expressed in the olfactory epithelium (OE), while Muc5ac is exclusively expressed in the nasal respiratory epithelium (Amini *et al*, 2019). The secreted protease from *E. coli* has mucin-binding activity through a 102 aa domain (designated X409 in Nason *et al*, 2021). X409 binds strongly to the O-glycosylated repeated mucin motifs, suggesting an interaction with the mucin TR backbone and innermost monosaccharide residues of attached O-glycans. Here, we designed a construct in which the antiviral C2 is fused to X409 to anchor it to the mucins, and we tested its efficiency in hamsters to limit SARS-CoV-2 infection when in contact with infected individuals.

## Results

To increase the residence time of the antiviral protein C2 in the nasal cavity, we produced C2X, a protein chimera made by C2 and the mucin-binding domain X409 separated by a flexible linker (**Supp. Fig. 1**). An overview of the strategy is shown in **Fig. 1A**. We first verified if the addition of the X409 domain to C2 interfered with its neutralizing property. Cell culture infection with D614G SARS-CoV-2 variant showed similar IC_50_ (6.5 nM and 13 nM for C2 and C2X, respectively; **Fig. 1B**) and we observed a better inhibition overall of SARS-CoV-2 pseudoviruses entry by C2X compared to C2 across different variants of concerns (VoCs), except for Beta VoC (**Fig. 1C**). We next assessed the effectiveness of the X409 mucin-binding domain to improve the residence time of C2 in the lumen of the nasal cavity of mice (**Fig. 1D**). While C2 was still observable 1h after nasal instillation in the mucus covering the OE, C2X was much more present at this time (**Fig. 1E, F**) and contrary to C2, was still found in the mucus layer 6h after nasal instillation.

**Figure 1:**
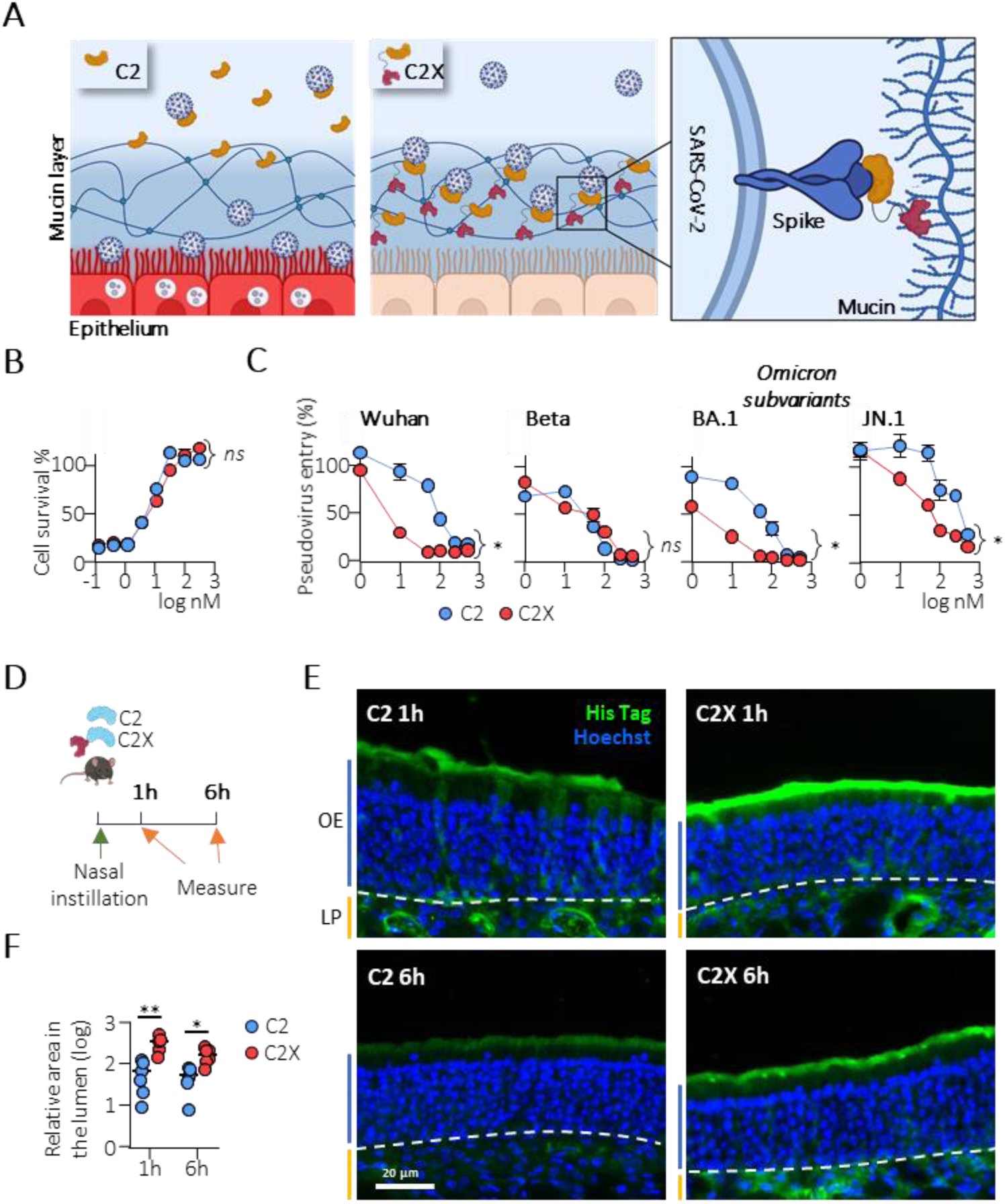
Validation of C2X antiviral activity and bioavailability in the nasal cavity. *(**A**) Strategy of coupling an antiviral protein (C2) with a mucin-binding domain (X) to improve its efficiency to block virus entry in vivo*, (**B**) Neutralisation *activity of C2X versus C2 against SARS-CoV-2 infection (D614G), (**C**) Entry blocking activity of C2X and C2 against SARS-CoV-2 variants of concerns pseudotypes. Data are presented as means ± SEM, n=3, two-way ANOVA; (ns not significant; *p<0.0001). (**D**) Experimental design to measure C2 and C2X residence time in mice after nasal instillation. (**E**) Representative images of C2 and C2X presence in the OE revealed by immunohistochemistry. (**F**) Measure of C2 and C2X area in the OE mucus layer. Data are presented as means ± SEM, n=6 nasal cavity from 3 mice, Mann-Whitney; (*p<0.05; **p<0.01)*.

We previously validated the ability of an improved version of C2 to decrease SARS-CoV-2 infection level in hamsters but observed only partial protection (Thébault *et al*, 2022). As targeting virus entry in the nasal cavity could be used to prevent infection during contact with contagious individuals, we evaluated the antiviral efficiency of C2X intranasal treatment after a 12h contact with a 1-day post-infection (dpi) hamster kept in the same cage as a PBS-treated animals used as a control sentinel of transmission for 60h (**Fig. 2A**). We choose a 1dpi hamster to transmit SARS-CoV-2 as the virus particle shedding is maximum at this stage (Merle-Nguyen *et al*, 2024). The treatment (PBS *vs* 60µg C2X) was started 1h before contact and repeated every 12h. Global pathophysiological impact of infection was first measured through weight loss which did not occur in C2X-treated animals (**Fig. 2B**). Using RT-qPCR to measure viral replication in the olfactory turbinates, we observed a transmission of the virus in all animals of the PBS-treated group while the olfactory turbinates were almost free of viral subgenomic RNA in 4 of the 6 C2X-treated animals. The 2 remaining animals present ∼1 log less level of viral sgRNA compared to PBS-treated animals (**Fig. 2C**). This difference was associated with a marked decrease in inflammatory cytokine expression levels in C2X-treated animals, similar to that seen in uninfected animals for TNF_α_ and IFN_λ_. We examined if these results would be consistent with immunohistochemistry performed on the same animals using the other side of the nasal cavity. In PBS-treated sentinel animals, we observed a typical infection of the OE along Iba1^+^ resident macrophages infiltration, epithelium damage and high level of cellular debris in the lumen of the nasal cavity (**Fig. 2E**). In C2X-treated sentinel animals, we observed a very low viral infection in the respiratory epithelium localized in the ventral part of the olfactory turbinates with an absence of epithelium damage and cellular debris in the lumen of the nasal cavity (**Fig. 2F**; **Supp. Fig. 2**). The level of Iba1^+^ cells infiltration in the OE was also much lower in C2X-treated sentinel animals compared to the PBS group and similar to the OE of uninfected animals. Consistent with a lower level of infection in the nasal cavity, we observed a decreased infection and inflammation in the lungs. While RT-qPCR revealed virus multiplication in the lungs of all animals of the PBS-treated group, the lungs were almost free of viral sgRNA in 3 among the 6 C2X-treated animals (**Fig. 2D**). The three others displayed ∼1 log lower level of viral subgenomic RNA compared to PBS-treated animals. This difference was reflected by lower inflammatory cytokines TNF_α_ and IFN_λ_ expression levels compared to untreated animals.

**Figure 2:**
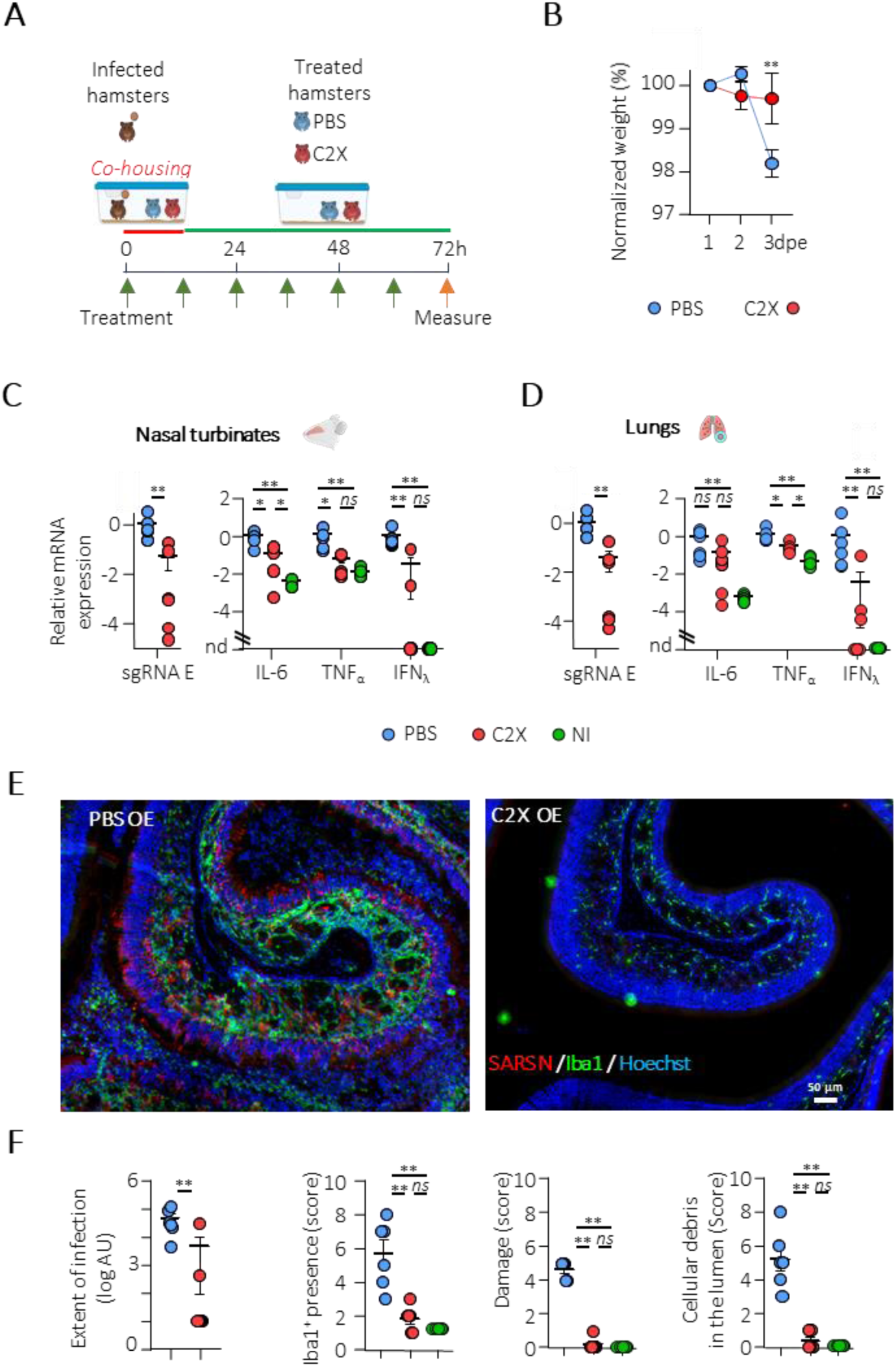
Effectiveness of C2X to limit airborne transmission of SARS-CoV-2 in hamsters. *(**A**) Experimental design of hamsters co-housing. A 1 dpi intranasally infected hamster was left for 12 hours in contact with individuals treated either with PBS (blue) or C2X every 12 hours until 3 days post exposition (dpe) (red). (**B**) Weight evolution following contact between infected animals and C2X-or PBS-treated hamsters. (**C, D**) Quantification of viral mRNA encoding SARS-CoV-2 E protein; IL-6, TNFa, IFNλ, in nasal turbinates and in the lungs, respectively. (**E**) Representative images of SARS-CoV-2 infection of the olfactory epithelium in C2X-treated animals revealing the SARS-CoV-2 N protein and the macrophage marker Iba1. (**F**) Measure of the area of infection; score of Iba1^+^ resident macrophages presence, damage as well presence of cellular debris in the lumen of the nasal cavity. AU, arbitrary unit; data are presented as means ± SEM, n=6, Mann-Whitney; (ns not significant; *p<0.05; **p<0.01)*.

As a chimeric protein constituted by an artificial protein (C2) and a bacterial mucin-binding domain (X409; Nason *et al*, 2021; Thébault *et al*, 2022), C2X could be inflammatory. To test it, mice were treated 3 days with C2X renewed 15 days apart (**Fig. 3A**). **Fig. 3B** shows that the levels of expression of inflammatory cytokines IFN_λ_, TNF_α_, IL1_β,_ IL17A, and of the chemokine CXCL10 as well as different markers linked to adaptative immune response were not different in C2X-treated animals compared to PBS-treated group. Similarly, we observed no difference for Iba1 and CD3e expression levels, respectively linked to macrophages and lymphocytes presence in the olfactory turbinates. These results are consistent with an absence of significant inflammatory effect of the repeated C2X treatments which was confirmed histologically by observing similar cellular density and activation state of Iba1^+^ resident macrophages (**Fig. 3C**). Furthermore, with an intranasal treatment meant to be used daily, C2X could trigger production of antibodies that block C2X interaction with its Spike target, making the treatment ineffective in the long term. We thus collected sera from the C2X-treated mice. While we observed a small decrease of C2X efficiency at the highest concentration of serum used (1/20), this was not different from the serum of mice treated with PBS indicating that the repeated treatment of C2X in mice did not produce blocking antibodies dampening its antiviral property (**Fig. 3D**). We finally observed an absence of C2X *in vivo* toxicity using zebrafish embryo acute toxicity test at concentration up to 50 times higher than what can be estimated from the dose used in hamsters and mice (**Fig. 3E**).

**Figure 3:**
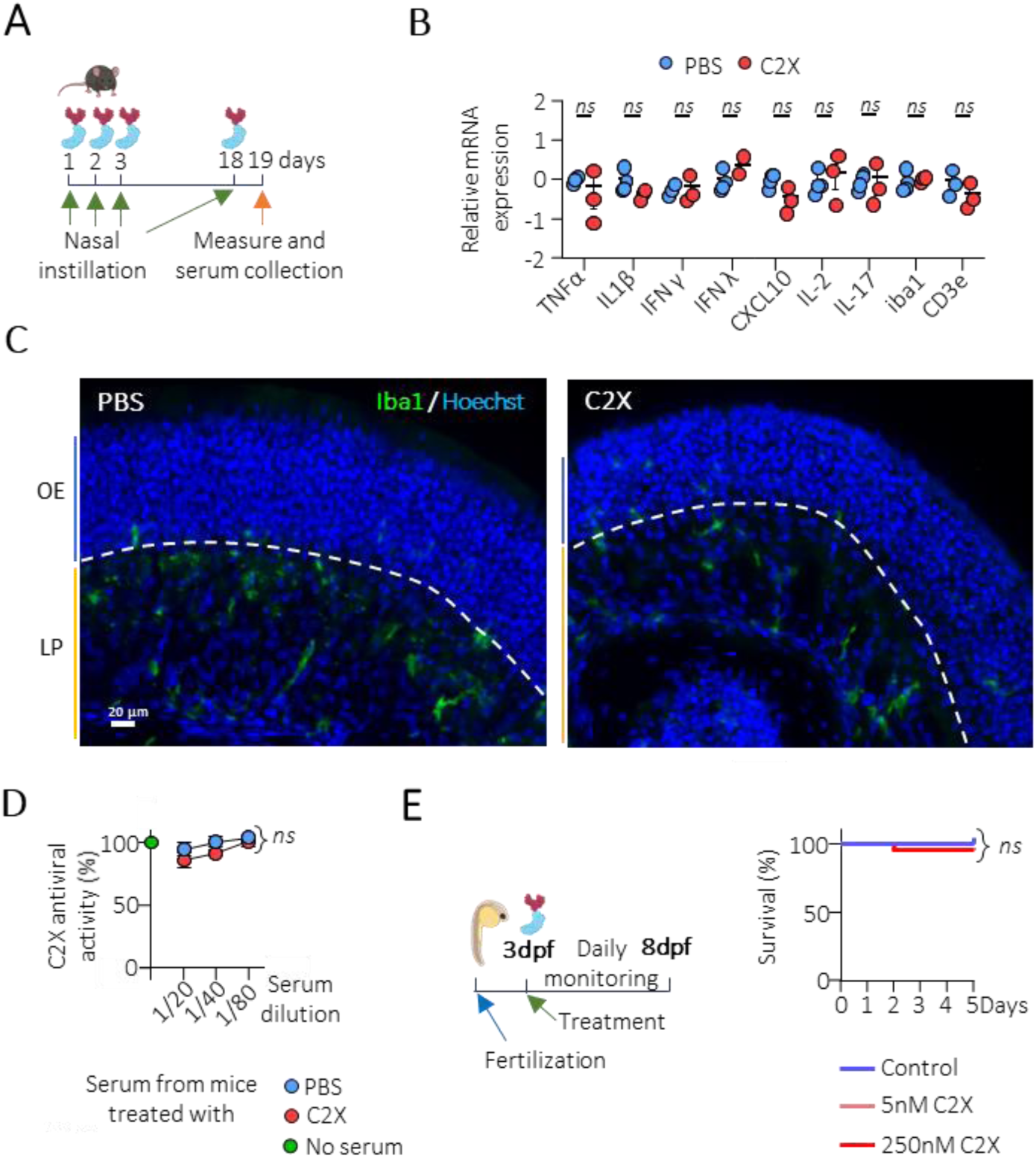
Absence of immunogenicity, inflammation and in vivo toxicity of C2X treatment. *(**A**) Experimental design to measure inflammation and antigenicity induced by C2X nasal instillation (**B**) Quantification of mRNA encoding inflammatory and immune markers, data are presented as means ± SEM, n=4, (Mann-Whitney; ns not significant) **(C)** Representative images of Iba1^+^ resident macrophages presence in the olfactory epithelium (OE) and the lamina propria (LP) of C2X-treated (C2X, n=3) and vehicle-treated (PBS, n=4) animals (**D**) Quantification of the anti-C2X activity of mice sera treated with C2X or PBS. Activity was measured as a percentage of inhibition of C2X on SARS-CoV-2 pseudotypes entry. Data are presented as means ± SEM, n=3, (two-way ANOVA; ns not significant). (**E**) Experimental design to measure toxicity of C2X in zebrafish embryo (left). Embryos were treated 3d post-fertilization with C2X at various concentrations and monitored daily during five days (n=20). Data are presented as a percentage of surviving embryos (right).*

## Discussion

In order to overcome the mucociliary clearance preventing an effective treatment of the nasal cavity, C2X was designed for mucus anchoring. Addition of the X409 mucin-binding domain results in an extended residence time of C2X *vs* C2 in the nasal cavity, with C2X present in the mucus layer 6h post-instillation. X409 has been reported to bind with high specificity Muc5ac which is one of the two airway polymeric mucins with Muc5b (Amini *et al*, 2019; Nason *et al*, 2021). To our knowledge, this is the first example of an antiviral nanobinder designed to be mucus-anchored to block virus infection. Using the hamster animal model of SARS-CoV-2 pathogenesis, we show that C2X, a version of a previously reported SARS-CoV-2 Spike neutralizing nanobinder, reduces SARS-CoV-2 airborne transmission and disease when administered to sentinel hamsters. We observed a decrease of more than 1 log average with 4 of the 6 C2X-treated animals almost spared from infection in the nasal turbinates (and 3 of the 6 in the lung tissues). We only observed traces of infection in the respiratory epithelium and Steno’s gland epithelium on the ventral part of the nasal cavity in C2X-treated animals (**Supp. Fig. 2**). In accordance with these results, inflammation markers of the infection were at very low levels in these animals. As these markers are the basis for the development of severe forms of COVID-19 in SARS-CoV-2-infected patients, it highlights the promising strategy to increase resident time of antivirals in the nasal cavity when in contact with infected individuals. Without the addition of the X409 mucin-binding domain, a daily treatment of an improved version of C2 used at higher concentration (600µg instead of 60µg) was barely effective to limit SARS-CoV-2 infection (reduction by a factor 2 of viral presence in the olfactory epithelium at 3 dpi; Thébault *et al*, 2022). These results show that the anchoring of neutralizing binders to the mucus is an efficient strategy to increase their *in vivo* antiviral activity and drastically limit the impact of viral infection-associated damage and inflammatory levels in infected tissues. This was emphasized by the very low increase of IFN_λ_ expression level in C2X-treated sentinels compared to PBS-treated one.

As the X409 mucin-binding domain and C2 backbone have both a procaryotic origin, the repeated treatment could be antigenic and lead to epithelium inflammation, potentially eliciting production of C2X neutralizing antibodies and epithelium damage, respectively. Our safety and toxicology studies did not reveal any adverse effect of repeated C2X treatment in mice and in zebrafish. Furthermore, C2X anchoring in the mucus did not appear to elicit an immune response able to interfere with its neutralizing activity.

The nasal cavity is now considered the primary target during respiratory viruses infection (Richard *et al*, 2020; Wellford & Moseman, 2023). These viruses can be transmitted very effectively with 50% SARS-CoV-2 infection rate with only 10 pfu in humans (Killingley *et al*, 2022) and a 4h contact in hamsters separated by 2 meters (Port *et al*, 2024). Our results show that the C2X treatment in the nasal cavity protects hamster sentinels in direct contact for 12h with infected animals at 1 dpi when the virus shedding is at the highest level (Merle-Nguyen *et al*, 2024). In our experimental setup, we kept the PBS and C2X treated animal for a following 60h, thus the PBS treated animals could in turn infect the C2X treated one. This very promising results could be improved by extending the avidity of C2X for both the Spike and the mucins by multimerization of C2X using dedicated sequence motifs such as the trimerizing foldon domain (Tao *et al*, 1997).

Overall, our data highlight the promise of antiviral nanobinders and their targeting to the mucus of the nasal cavity as potent viral strategy to block respiratory virus infection when in contact with infected individuals. This strategy could be extended easily to other respiratory viruses such as influenza and respiratory syncytial virus with specific entry blocking nanobinders (Rossey *et al*, 2017).

## Material & Methods

### Ethics statement

The study was carried out following protocols approved by INRAE and ANSES/ENVA/UPEC local Ethics Committee for mice and hamsters experiments respectively. Both were authorized by the French Ministry of Research under the number APAFIS #49851-2023112816322635v4 and APAFIS #35556-2022022322445750 v7, respectively, in accordance with French and European regulations.

### Antiviral proteins expression and purification

C2 and C2X open reading frames were subcloned on pQE81L bacterial expression vector to drive their expression in *E. coli* as His-tagged proteins. Gene expression was induced by addition of IPTG overnight at 37°C. Bacteria were pelleted and resuspended in 200mM NaCl, 20mM Tris pH7.4, containing a cocktail of protease inhibitors (Roche Diagnostics GmbH). Bacteria were sonicated on ice and lysates clarified by centrifugation at 10,000g for 30 min at 4°C. Supernatants were submitted to affinity chromatography on HisTrap columns (GE Healthcare Life Sciences). Fractions containing the protein of interest were pooled and injected in a gel filtration Superdex S200 column equilibrated with PBS. Fractions containing purified proteins were identified by SDS-PAGE and frozen at -80°C.

### Virus stock production

For *in vitro* experiments, SARS-CoV-2 isolate France/IDF0372/2020 was kindly provided by Sylvie van der Werf. It was propagated in Vero-E6 cells in Dulbecco’s modified Eagle’s medium (DMEM) supplemented with 2% (v/v) foetal bovine serum (FBS, Invitrogen). Viral titres were determined by plaque assay in Vero-E6 cells (CRL-1586, ATCC).

### Neutralization assay

To measure neutralization activity, Vero-E6 cells were seeded in 96-well plates, 2 x 10^4^/well, in Dulbecco’s modified Eagle’s medium (DMEM) supplemented with 10% (v/v) foetal bovine serum (FBS, Invitrogen) and incubated overnight at 37°C. One day later, dilutions of C2 and C2X were mixed with an equal volume of medium containing 2 x 10^3^ PFU (corresponding to an multiplicity of infection of 0.1) of virus in DMEM for 1h at room temperature before addition and 72h incubation at 37°C. Cell viability was measured by CellTiter-Glo reagent (CellTiter-Glo Luminescent Cell Viability Assay, Promega #G7571). Luminescence was quantified using the Infinite M200Pro TECAN apparatus.

### Pseudotypes entry blocking assay

#### Pseudotype production

Murine leukaemia virus pseudo-typed particles expressing the Spike protein of different variants of SARS-CoV-2 were produced as previously described (Thebault et al., 2022). Four different pseudotypes were produced, a first containing the Spike of SARS-CoV-2 D614G (Genbank accession number: MN908947); and the three others containing the Spike with mutations within the receptor-binding domain representatives of the β variant (N501Y, K417N and E484K mutations), the Omicron BA.1 variant (S371L, S373P, S375F, K417N, N440K, G446S, S477N, T478K, E484A, Q493K, G496S, Q498R, N501Y and Y505H substitutions); and the Omicron JN.1 variant (with the complete substitution of the amino acid domain [1-516] by its homolog defined by the Genebank number WXB27687 sequence).

#### Entry blocking assay

HEK-293T-hACE2 cells were seeded in a 96-well plate, 2 x 10^4^/well, in DMEM complemented with 10% FBS (v/v) and incubated at 37°C. At day 1, C2 and C2X were mixed with pseudo-typed particles for 1h at room temperature before addition and 48h incubation at 37°C. Cells were lysed as indicated in Promega # E1501 Luciferase Assay system protocol and luciferase activity was measured using the Infinite M200Pro TECAN apparatus.

### Animal experiments

For C2 vs C2X residence time in the nasal cavity, two groups of 3 Balb/c mice were intranasally instilled with PBS (pH 7.4) containing C2 or C2X (60 µg). Animals were euthanised 1 h or 6 h later for tissue collection.

For C2X immunogenicity and immunity impact; two groups of 3 and 4 Balb/c mice were treated intranasally once a day with 60 µg of C2X or PBS respectively for 3 consecutive days. The treated mice were kept for 15 days. One day before euthanasia, mice were treated with an additional dose of C2X or PBS. Nasal cavity was collected and divided into two halves: one for histology and immunofluorescence assays and a second for RT-qPCR analysis. Sera were collected and conserved at -80°C.

For hamster infection challenge, 24 specific-pathogen-free (SPF) 8-week-old male Golden Syrian hamsters *Mesocricetus auratus* (Janvier-Labs, Le Genest-Saint-Isle, France) housed under BSL-III conditions were kept according to the standards of French law for animal experimentation. Six hamsters were used as control. Six hamsters were infected with 5.10^3^ TCID_50_ of SARS-CoV-2 isolate FRANCE/IDF0372/2020 one day before co-housing started with two sentinel hamsters. Sentinels were treated by nasal instillation with 80µL of PBS or C2X (60 µg of protein/hamster) 1h before the 12h co-housing. Infected hamsters were separated from sentinels and treatment continued twice a day before being euthanized at 3 dpe (days post exposition). The head of each sentinel was collected, it was separated into two hemi-heads, one half was used for histology and the second for RT-qPCR analysis.

### mRNA quantification (RT-qPCR)

Total RNA was extracted from frozen nasal turbinates and lungs using the Trizol-chloroform method as previously described (Bourgon *et al*, 2022) with some modifications. Briefly, Oligo-dT first strand mixed with random hexamers cDNA synthesis was performed from 1 μg total RNA by iScriptAdv cDNA kit (Biorad, #1725038) following the manufacturers recommendations. qPCR was carried out using 125 ng of cDNA templates added to a 15 μL reaction mixture containing 500 nM primers (sequences in **Table 1**) and iTaq Universal SYBR Green Supermix (Biorad, # 1725124) using a thermocycler (Mastercycler ep realplex, Eppendorf). Fluorescence was monitored and measured by Realplex Eppendorf Software. A dissociation curve was plotted at the end of the 40 amplification cycles of the qPCR to check the ability of theses primers to amplify a single and specific PCR product. Quantification of initial specific RNA concentration was done using the ΔΔCt method (Muller *et al*, 2002). Standard controls of specificity and efficiency of the qPCR were performed. The mRNA expression of each gene was normalized with the mean expression level of dicer/β-actin genes.

### Immunohistochemistry

Mouse and hamster nasal tissues were fixed in 4% Neutral Buffer Formalin (F0043, Diapath) for 1 or 3 days at 4°C respectively, then decalcified in Osteosoft (1017281000, Merck) at 4°C for 1 or 3 weeks, respectively. The nasal cavities were cryoprotected with sucrose (30%) and embedded with Epredia Neg-50 (11912365, FisherScientific). Cryosectioning was performed on median transversal sections, perpendicular to the hard palate, highlighting the nasal structures and epithelia. Non-specific staining was blocked by incubation with 2% bovine serum albumin (BSA) and 0.05% Tween. The sections were then incubated overnight with primary antibodies directed against His-tag (Mouse; 1:500; MA1-21315; Invitrogen); SARS-CoV-2 N (Mouse 1:1000; clone 1C7C7; Sigma Aldrich); ionized calcium-binding adapter molecule 1 (Iba1) (Rabbit; 1:800; Molecular Probes–A11001; Invitrogen). Fluorescence staining was performed using goat anti-mouse-A555; goat anti-rabbit-A488 (Molecular Probes A21422; A11056 respectively). Images were taken with an Olympus IX83 inverted microscope equipped with an Orca-Fusion (Hamamatsu, C14440-20UP). Infection levels were measured by quantification of SARS-N fluorescence area with ImageJ (Rasband, W.S., ImageJ, U. S. National Institutes of Health, Bethesda, Maryland, USA, http://imagej.nih.gov/ij/, 1997–2012). Level of epithelium damage and cellular debris in the lumen were scored from 0 (absence of damage and cellular debris) to 10 (fully desquamated epithelium and lumen filled with cellular debris) based on DAPI staining as performed previously (Bourgon et al, 2022). Iba1^+^ staining was used to scored resident macrophages infiltration from 0 (absence of staining) to 10 (massive infiltration of Iba1^+^ cells). All quantifications were performed blind to the treatment.

### Toxicity assessment

Wild-type (AB strain) zebrafish (*Danio rerio*) eggs were fecundated and incubated at 28°C in E3 medium (5 mM NaCl, 0.17 mM KCl, 0.33 mM CaCl2, 0.33 mM MgSO4) supplemented with 0.3 µg/mL methylene blue. Three days after fecundation, the embryos were incubated in water with 5 nM or 250 nM of C2X during 5 days (n=20 for each group). Apical observations of the fish were made daily to detect indicators of lethality (coagulation of fertilised eggs, lack of somite formation, lack of detachment of tail-bud from yolk sac, lack of heartbeat and mortality as well as any sign of morbidity).

### Statistical analysis

All statistical comparisons were performed using Prism 8 (GraphPad). Data shown as the means ± SEM. Quantitative data were compared across groups using two-way ANOVA test for neutralization assay with SARS-CoV-2 pseudovirus entry assays and weight measures. All other parameters were tested using Mann-Whitney. Detailed information on statistical test used, sample size and p-value are provided in the Figure legends.

## Acknowledgements

NM is supported by INRAe SA department; ANRS (Grant UCRAH). We thank Christelle Langevin and Marion Mehraz from IERP (UE 0907, Infectiologie Expérimentale des Rongeurs et Poissons) for experimentation on zebrafish; Julie Rivière and Martha Vilotte from @BRIDGe (UMR1313 Génétique Animale et Biologie Intégrative) for their collaboration in the histology scanning; Bruno da Costa (UR892, Unité de Virologie et Immunologie Moléculaires) for the production of live virus used in the neutralization assay and the PRBM platform of ENVA for their helped in the BSL3 animal facility and Birte Nielsen for her help improving the manuscript.

## Authors contribution

Conceptualization: NM and BD. Investigation: JGR, AD, JJ, NL, QN, BK, SLP, BD and NM. Writing: NM and BD, with input from all authors.

---

**Supplementary figure 1:**
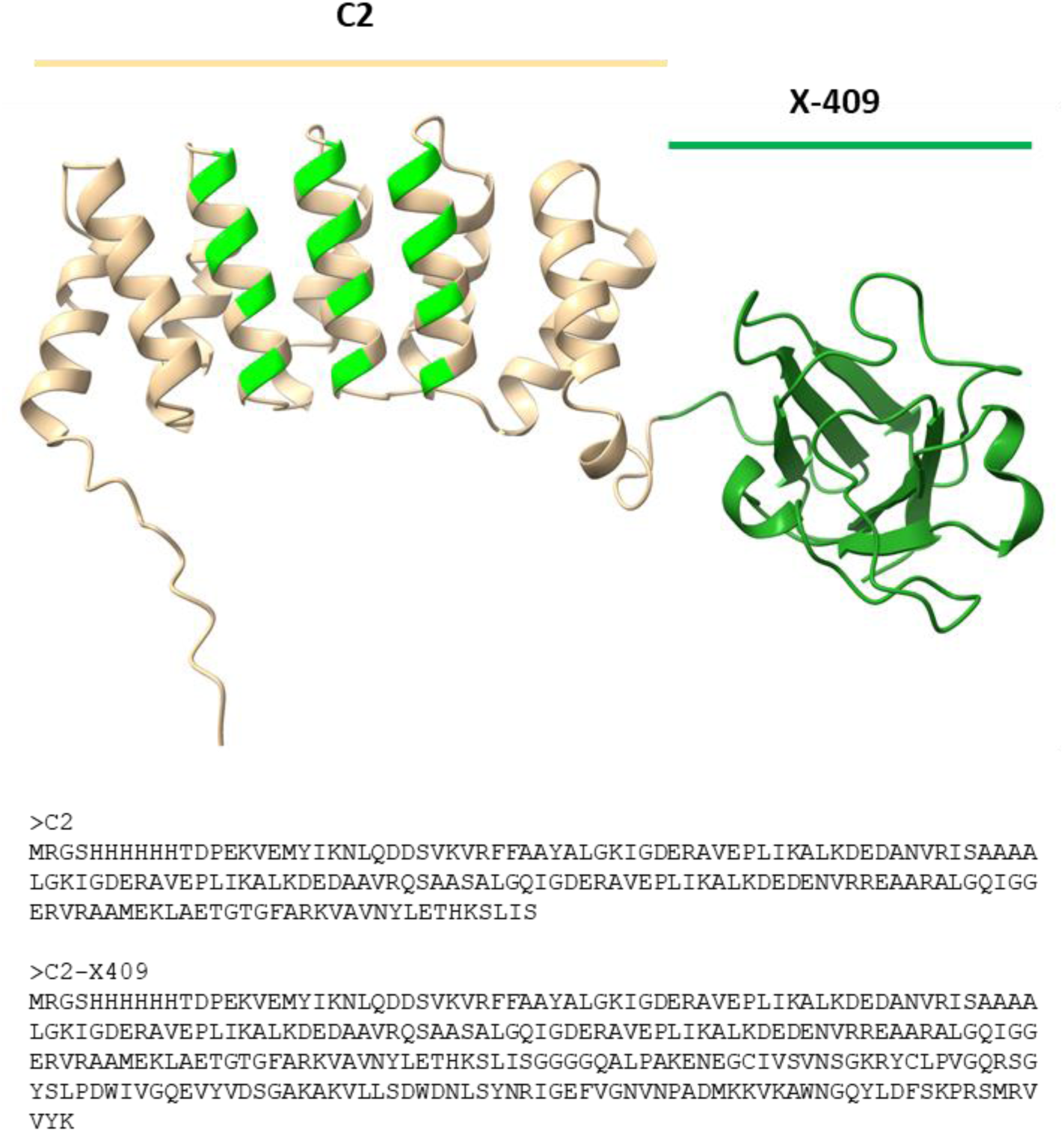
Structure and sequences of C2 and C2X (Structure predicted using AlphaFold 3)

**Supplementary figure 2:**
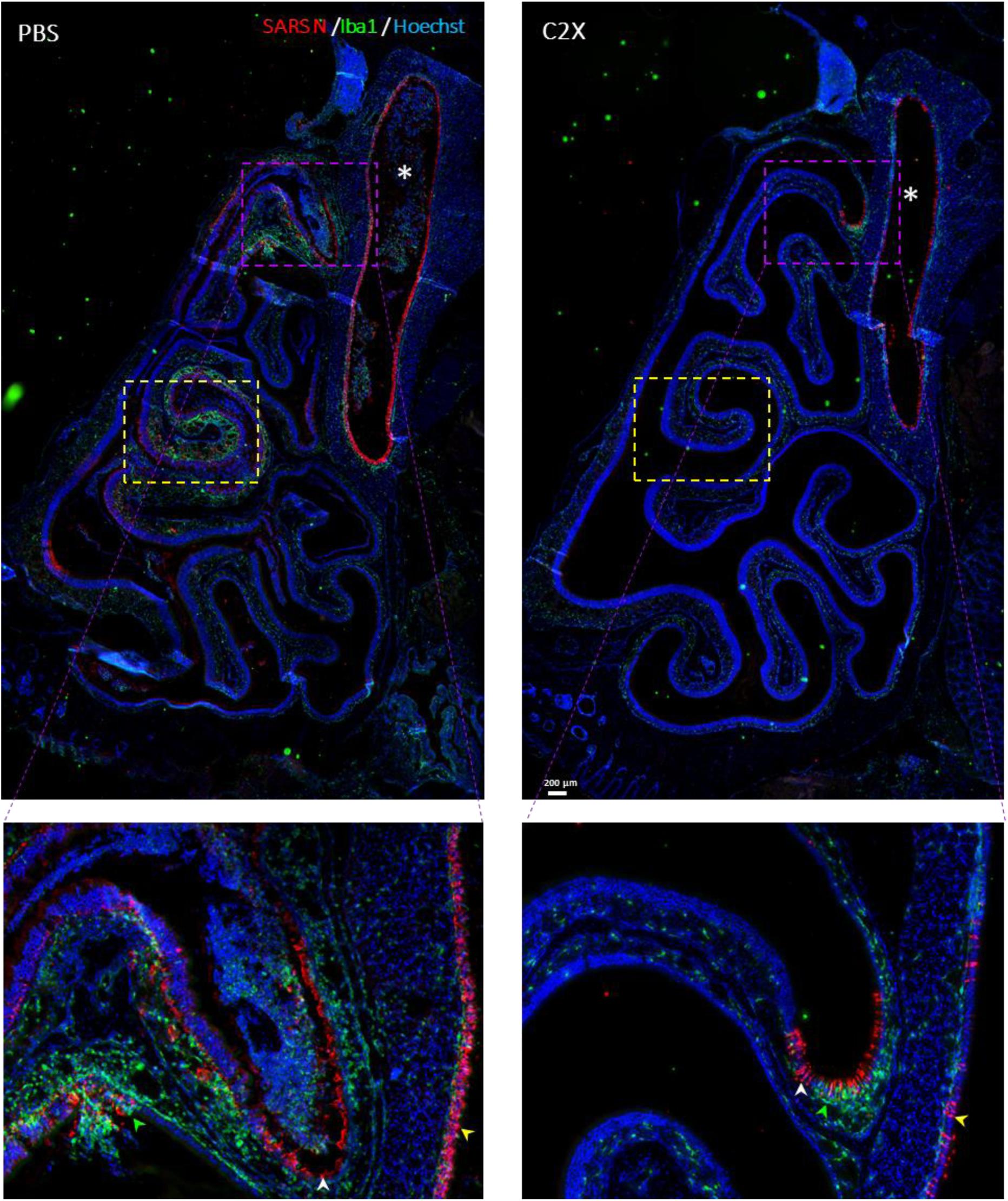
Representative images of posterior zone of the nasal cavity containing mainly olfactory epithelium from hamster sentinels at 3 dpe (days post exposition). Magnified area (dashed violet square) showing presence of infected cells in respiratory epithelium (white arrow heads); Steno’s gland epithelium (yellow arrow heads) as well as Iba1^+^ resident macrophages infiltration (green arrow heads). This area is the same as used in *Figure 2* (yellow dashed square).

## References

1. Amini SE, Gouyer V, Portal C, Gottrand F & Desseyn JL (2019) Muc5b is mainly expressed and sialylated in the nasal olfactory epithelium whereas Muc5ac is exclusively expressed and fucosylated in the nasal respiratory epithelium. Histochemistry and cell biology

2. Bryche B, St Albin A, Murri S, Lacôte S, Pulido C, Ar Gouilh M, Lesellier S, Servat A, Wasniewski M, Picard-Meyer E, et al (2020) Massive transient damage of the olfactory epithelium associated with infection of sustentacular cells by SARS-CoV-2 in golden Syrian hamsters. Brain Behav Immun 89: 579–586

3. Chen TM & Dulfano MJ (1978) Mucus viscoelasticity and mucociliary transport rate. J Lab Clin Med 91: 423–431

4. Corfield AP (2015) Mucins: A biologically relevant glycan barrier in mucosal protection. Biochimica et Biophysica Acta (BBA) - General Subjects 1850: 236–252

5. Gizurarson S (2015) The Effect of Cilia and the Mucociliary Clearance on Successful Drug Delivery. Biological & Pharmaceutical Bulletin 38: 497–506

6. Hou YJ, Okuda K, Edwards CE, Martinez DR, Asakura T, Dinnon KH, Kato T, Lee RE, Yount BL, Mascenik TM, et al (2020) SARS-CoV-2 Reverse Genetics Reveals a Variable Infection Gradient in the Respiratory Tract. Cell 182: 429–446.e14

7. Killingley B, Mann AJ, Kalinova M, Boyers A, Goonawardane N, Zhou J, Lindsell K, Hare SS, Brown J, Frise R, et al (2022) Safety, tolerability and viral kinetics during SARS-CoV-2 human challenge in young adults. Nat Med 28: 1031–1041

8. Kozlov M (2022) Could a nose spray a day keep COVID away? Nature: d41586–022-03341-z

9. Marttin E, Schipper NGM, Verhoef JC & Merkus FWHM (1998) Nasal mucociliary clearance as a factor in nasal drug delivery. Advanced Drug Delivery Reviews 29: 13–38

10. Mayor A, Thibert B, Huille S, Respaud R, Audat H & Heuzé-Vourc’h N (2021) Inhaled antibodies: formulations require specific development to overcome instability due to nebulization. Drug Deliv Transl Res 11: 1625–1633

11. Merle-Nguyen L, Ando-Grard O, Bourgon C, St Albin A, Jacquelin J, Klonjkowski B, Le Poder S & Meunier N (2024) Early corticosteroid treatment enhances recovery from SARS-CoV-2 induced loss of smell in hamster. Brain, Behavior, and Immunity 118: 78–89

12. Muller PY, Janovjak H, Miserez AR & Dobbie Z (2002) Processing of gene expression data generated by quantitative real-time RT-PCR. Biotechniques 32: 1372–1374, 1376, 1378–1379

13. Nason R, Büll C, Konstantinidi A, Sun L, Ye Z, Halim A, Du W, Sørensen DM, Durbesson F, Furukawa S, et al (2021) Display of the human mucinome with defined O-glycans by gene engineered cells. Nat Commun 12: 4070

14. Port JR, Morris DH, Riopelle JC, Yinda CK, Avanzato VA, Holbrook MG, Bushmaker T, Schulz JE, Saturday TA, Barbian K, et al (2024) Host and viral determinants of airborne transmission of SARS-CoV-2 in the Syrian hamster. eLife 12: RP87094

15. Richard M, van den Brand JMA, Bestebroer TM, Lexmond P, de Meulder D, Fouchier RAM, Lowen AC & Herfst S (2020) Influenza A viruses are transmitted via the air from the nasal respiratory epithelium of ferrets. Nat Commun 11: 766

16. Rossey I, Gilman MSA, Kabeche SC, Sedeyn K, Wrapp D, Kanekiyo M, Chen M, Mas V, Spitaels J, Melero JA, et al (2017) Potent single-domain antibodies that arrest respiratory syncytial virus fusion protein in its prefusion state. Nat Commun 8: 14158

17. Tao Y, Strelkov SV, Mesyanzhinov VV & Rossmann MG (1997) Structure of bacteriophage T4 fibritin: a segmented coiled coil and the role of the C-terminal domain. Structure 5: 789–798

18. Thébault S, Lejal N, Dogliani A, Donchet A, Urvoas A, Valerio-Lepiniec M, Lavie M, Baronti C, Touret F, Da Costa B, et al (2022) Biosynthetic proteins targeting the SARS-CoV-2 spike as anti-virals. PLoS Pathog 18: e1010799

19. Valdez-Cruz NA, García-Hernández E, Espitia C, Cobos-Marín L, Altamirano C, Bando-Campos CG, Cofas-Vargas LF, Coronado-Aceves EW, González-Hernández RA, Hernández-Peralta P, et al (2021) Integrative overview of antibodies against SARS-CoV-2 and their possible applications in COVID-19 prophylaxis and treatment. Microb Cell Fact 20: 88

20. Wellford SA & Moseman EA (2023) Olfactory immunology: the missing piece in airway and CNS defence. Nat Rev Immunol

